# *Wolbachia* in the spittlebug *Prosapia ignipectus:* Variable infection frequencies, but no apparent effect on host reproductive isolation

**DOI:** 10.1101/2021.02.25.432892

**Authors:** Timothy B. Wheeler, Vinton Thompson, William R. Conner, Brandon S. Cooper

## Abstract

Animals serve as hosts for complex communities of microorganisms, including endosymbionts that live inside their cells. *Wolbachia* bacteria are perhaps the most common endosymbionts, manipulating host reproduction to propagate. Many *Wolbachia* cause intense cytoplasmic incompatibility (CI) that promotes their spread to high and relatively stable frequencies. *Wolbachia* that cause weak or no CI tend to persist at intermediate, often variable, frequencies. *Wolbachia* could also contribute to host reproductive isolation (RI), although current support for such contributions is limited to a few systems. To test for *Wolbachia* frequency variation and effects on host RI, we sampled several local *Prosapia ignipectus* (Fitch)(Hemiptera: Cercopidae) spittlebug populations in the northeastern USA over two years, including closely juxtaposed Maine populations with different monomorphic color forms, “black” and “lined”. We discovered a group-B *Wolbachia* (*w*Pig) infecting *P. ignipectus* that diverged from group-A *Wolbachia—like* model *w*Mel and *w*Ri strains in *Drosophila—6* to 46 MYA. Populations of the sister species *Prosapia bicincta* (Say) from Hawaii and Florida are uninfected, suggesting that *P. ignipectus* acquired *w*Pig after their initial divergence. *w*Pig frequencies were generally high and variable among sites and between years. While phenotyping *w*Pig effects on host reproduction is not currently feasible, the *w*Pig genome contains three divergent sets of CI loci, consistent with high *w*Pig frequencies. Finally, Maine monomorphic black and monomorphic lined populations of *P. ignipectus* share both *w*Pig and mtDNA haplotypes, implying no apparent effect of *w*Pig on the maintenance of this morphological contact zone. We hypothesize *P. ignipectus* acquired *w*Pig horizontally as observed for many *Drosophila* species, and that significant CI and variable transmission produce high but variable *w*Pig frequencies.

## Introduction

Animals interact with microorganisms that influence their behavior, physiology, and fitness (Hurst and Jiggins, 2000; Brownlie *et al*., 2009; McFall-Ngai *et al*., 2013; Fredericksen *et al*., 2017; Gould *et al*., 2018; Hague, Caldwell and Cooper, 2020). These include associations between hosts and vertically transmitted endosymbionts that live inside their cells (McCutcheon, Boyd and Dale, 2019). Hosts may acquire endosymbionts cladogenically from common ancestors (Raychoudhury *et al*., 2009; Koga *et al*., 2013; Toju *et al*., 2013), from sister species via hybridization and introgression (Turelli *et al*., 2018; Cooper *et al*., 2019), or horizontally in ways that are not fully understood (O’Neill *et al*., 1992; Huigens *et al*., 2000; Ahmed *et al*., 2015). While few examples exist, endosymbionts can contribute to host reproductive isolation (RI) and speciation (Coyne and Orr, 2004; Matute and Cooper, 2021), highlighting the importance of discovering and characterizing endosymbiont-host associations.

Maternally transmitted *Wolbachia* bacteria are widely distributed (Werren, Baldo and Clark, 2008; Zug and Hammerstein, 2012; Weinert *et al*., 2015), infecting many arthropods and two groups of parasitic nematodes (Bandi *et al*., 1998), making *Wolbachia* perhaps the most common endosymbiont in nature. In *Drosophila,* introgressive and horizontal *Wolbachia* acquisition seem to predominate (Conner *et al*., 2017; Turelli *et al*., 2018; Cooper *et al*., 2019), but cladogenic acquisition during host speciation has been observed in other taxa (Raychoudhury *et al*., 2009; Gerth and Bleidorn, 2017). Many *Wolbachia* manipulate host reproduction to propagate in host populations. For example, many strains cause cytoplasmic incompatibility (CI) that reduces the egg hatch of uninfected embryos fertilized by *Wolbachia*-infected sperm (Hoffmann and Turelli, 1997). However, if females are also infected, the embryos survive, “rescuing” CI and promoting *Wolbachia* spread to high frequencies (Hoffmann, Turelli and Harshman, 1990; Turelli and Hoffmann, 1995; Barton and Turelli, 2011; Kriesner *et al*., 2013).

*Wolbachia* may contribute to host RI (Coyne and Orr, 2004; Matute and Cooper, 2021), with the best evidence coming from *Drosophila*. *Wolbachia* contribute to assortative mating and postzygotic isolation between co-occurring *D. paulistorum* semi-species (Miller, Ehrman and Schneider, 2010), and to reinforcement of isolation between uninfected *D. subquinaria* and *Wolbachia-infected D. recens* (Shoemaker, Katju and Jaenike, 1999; Jaenike *et al*., 2006). In contrast, *Wolbachia* do not contribute to RI in the *D. yakuba* clade, which includes *w*Yak- infected *D. yakuba, w*San-infected *D. santomea,* and *w*Tei-infected *D. teissieri* (Cooper *et al*., 2017). Thus, while some results from *Drosophila* strongly support contributions of *Wolbachia* to RI, and interest in the possibility of such effects remains high, it is unknown whether *Wolbachia* effects on RI are common in nature.

*Wolbachia* frequencies differ significantly among infected host taxa, ranging from very low to obligately fixed infections (Bandi *et al*., 1998; Kriesner *et al*., 2013; Cooper *et al*., 2017; Miller, Ehrman and Schneider, 2010). *Wolbachia* effects on reproduction (e.g., CI) and fitness, in combination with imperfect maternal transmission, govern its frequencies in host populations (Caspari and Watson, 1959; Hoffmann, Turelli and Harshman, 1990). Intensive sampling of a few systems has revealed both stable and variable *Wolbachia* frequencies within host populations. *Wolbachia* that cause intense CI, like *w*Ri in *Drosophila simulans,* persist at high and relatively stable frequencies, balanced by imperfect maternal transmission (Kriesner *et al.*, 2013; Turelli *et al*., 2018). In contrast, *Wolbachia* that cause weak or no CI tend to occur at variable intermediate frequencies (Hoffmann, Clancy and Duncan, 1996; Hamm *et al*., 2014; Kriesner *et al*., 2016; Cooper *et al*., 2017; Meany *et al*., 2019). These include *w*Mel-like *Wolbachia* frequencies that vary spatially in *D. melanogaster* and *D. yakuba* (Kriesner *et al.*, 2016; Hague *et al*., 2020), and temporally in *D. yakuba* and *D. santomea* (Cooper *et al*., 2017; Hague, Caldwell and Cooper, 2020). In all but a few systems, limited sampling has left a gap in knowledge about whether *Wolbachia* frequency variation is common (Hughes *et al*., 2011; Hamm *et al*., 2014; Cattel *et al*., 2016; Schuler *et al*., 2016; Ross *et al*., 2020).

*Prosapia ignipectus* (Fitch) (Hemiptera: Cercopidae) is one of about 14 species of *Prosapia* and one of two commonly found in the USA, the other being its sister species *P. bicincta* (Say)(Hamilton, 1977). *P. ignipectus* occurs in southern Ontario, Canada and the northeastern USA from Minnesota to Maine (Hamilton, 1977, 1982; Peck, 1999; Carvalho and Webb, 2005; Thompson and Carvalho, 2016). These species vary in male genital morphology and in associations with host plants, with *P. ignipectus* monophagous on the late season C4 perennial grass *Schizachyrium scoparium* (Little bluestem) (Hamilton, 1982; Thompson, 2004) and *P. bicincta* polyphagous on a variety of C4 grasses, but not including Little bluestem (Fagan and Kuitert, 1969; Thompson, 2004). Both species have conspicuous dorsal coloration, standing out against their respective host plants. All *P. bicincta* individuals have a single narrow transverse orange line across the widest part of the pronotum and a pair of narrow orange lines across the elytra. Most *P. ignipectus* individuals have a solid black dorsal surface, but in Maine some *P. ignipectus* have *P. bicincta*-like coloration (Figure 1). Notably, only 10 km separate monomorphic black and monomorphic lined *P. ignipectus* populations in western Maine, with little evidence of a hybrid zone and no obvious physical barriers to mixing across the boundary (Thompson and Carvalho, 2016). This morphological contact zone has persisted for at least 90 years. About 45 km southwest of this abrupt transition between aposematic color forms, three other *P. ignipectus* populations were found to be polymorphic with both black and lined forms— these populations are surrounded by monomorphic black populations. It has been hypothesized that *Wolbachia* determined RI may contribute to preservation of the sharp Maine morphological contact zone (Thompson and Carvalho, 2016).

**Figure 1.**
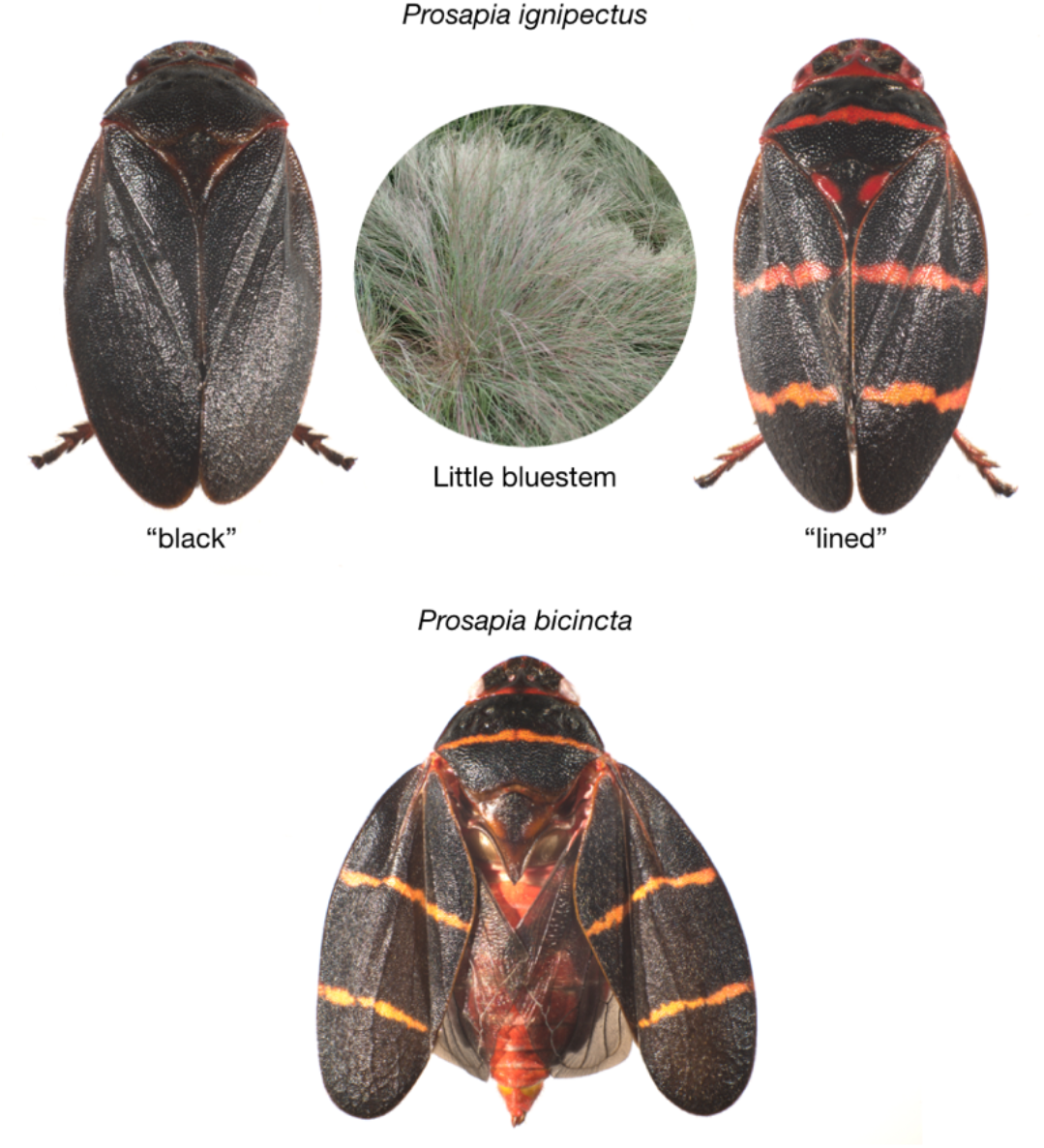
Sister species *P. ignipectus* and *P. bicincta* have conspicuous dorsal coloration. All *P. bicincta* individuals have a single narrow transverse orange line across the widest part of the pronotum and a pair of narrow orange lines across the elytra. Most *P. ignipectus* individuals have a solid black dorsal surface, but in Maine some *P. ignipectus* have *P. bicincta-like* coloration. *P. ignipectus* monophagous on the late season C4 perennial grass *Schizachyrium scoparium* (Little bluestem). Little bluestem photo by Krzysztof Ziarnek, Kenraiz (CC BY-SA 4.0, https://creativecommons.org/licenses/by-sa/4.0).

Here, we use collections of *P. ignipectus* from several sites in the northeastern USA across two years, in combination with collections of *P. bicincta* from Hawaii and Florida, USA, to assess modes of *Wolbachia* acquisition and to test for *Wolbachia* frequency variation through space and time. By sampling monomorphic black and lined populations and typing both *Wolbachia* and mtDNA haplotypes, we also test for contributions of *Wolbachia* to host RI. Finally, we generate whole genome *Wolbachia* data for phylogenetic analysis and to search for loci associated with inducing and rescuing CI (Beckmann, Ronau and Hochstrasser, 2017; LePage *et al*., 2017; Shropshire *et al*., 2018).

## Methods

### Sampling

We netted specimens from Little bluestem; sorted them by species, sex, and color form; and preserved them in 95% ethanol. The 2019 specimens *(N* = 4 sites) were collected on August 23. The 2020 specimens (*N* = 9 sites) were collected on August 9 (Silver Lake, NH), August 17 (Wonalancet, NH), and August 20 (all Maine localities) (Supplemental Table 1). Collection sites were on the verges of public rights of way or privately owned land. In two cases (New Vineyard and New Portland) they correspond to sites reported in Thompson and Carvalho (2016). Specimens were collected near the height of abundance for *P. ignipectus*, which starts to emerge in adult form in late July and early August. We also sampled three additional spittlebug species at these sites: *Lepyronia quadrangularis* (Say) (*N* = 25), *Philaenus spumarius* (L.) (*N* = 5), and *Philaenarcys killa* (Hamilton)(*N* = 24), all of the family Aphrophoridae. Like, *P. ignipectus, P. killa* a is monophage on Little bluestem. *L. quadrangularis* is a polyphage but often abundant on Little bluestem. *P. spumarius* is an extreme polyphage, with a preference for forbes (herbaceous perennial dicots) but is occasionally collected from Little bluestem in the company of *P. ignipectus*. By screening them for *Wolbachia* we tested for the possibility of horizontal *Wolbachia* transfer through plant interactions (Chrostek *et al.*, 2017). Lastly, because identification of infections in sister hosts enables formal analysis of modes of *Wolbachia* acquisition (Turelli *et al*., 2018; Conner *et al*., 2017; Cooper *et al*., 2019; Raychoudhury *et al*., 2009), we also obtained samples of the sister species *P. bicincta* from Hawaii (*N* = 60) and Florida (*N* = 40) to screen for infections. *P. bicincta* is native to the southeastern USA (Fagan and Kuitert, 1969; Thompson and Carvalho, 2016), but has recently been introduced into the Kona Region of Big Island, Hawaii (Thorne *et al*., 2018).

### *Wolbachia* typing

We generated whole genome *Wolbachia* data to type the *Wolbachia* infecting *P. ignipectus* and to search for loci associated with CI. We extracted 800ng of high molecular weight DNA (Qiagen Genomic-tip 20/G; Qiagen, Germany) from one black New Vineyard female (see below), and then input and sequenced it (Ligation Sequencing Kit, SQK-LSK109; FLO-MIN106 flow cell) for 48 hours (Oxford Nanopore Technologies). We mapped raw nanopore reads (5.8Gb of data) to all known *Wolbachia* sequences (NCBI taxid 953) with BLASTn and extracted reads where at least 60% of their length mapped (qcovs >= 60). We then corrected and assembled reads using canu 2.1.1 (Koren *et al*., 2017, 2018; Nurk *et al*., 2020) and polished the *Wolbachia* assembly using nanopolish 0.13.2 (Loman, Quick and Simpson, 2015). We annotated our *Wolbachia* assembly plus the genomes of model group-A (*w*Mel, Wu *et al*., 2004; and *w*Ri, Klasson *et al*., 2009) and group-B (*w*Pip-Pel, Klasson *et al*., 2008; and *w*Mau, Meany *et al*., 2019) strains using Prokka v.1.11 (Seemann, 2014). We used only genes present in single copy and with identical lengths in all genomes. To assess the quality of our assembly, we excluded *w*Pig and repeated this with only *w*Mel, *w*Ri, *w*Pip, and *w*Mau.

Preliminary analysis of a few loci placed the *P. ignipectus Wolbachia* in group-B (see below), but we performed Bayesian analyses using the GTR + Γ + I model for sequence evolution using whole genome data to confirm this (Höhna *et al*., 2016). Genes were concatenated and partitioned by codon position, with a rate multiplier, σ, assigned to each partition to accommodate variable substitution rates. We used flat, symmetrical Dirichlet priors on the stationary base frequencies, π, and the relative-rate parameters, η, of the GTR model (i.e., Dirichlet(1,1,1...)). As in Turelli et al. (2018), we used a Γ(2,1) hyperprior on the shape parameter, α, of the discrete-Γ model (adopting the conventional assumption that the β rate parameter equals α, so that the mean rate is 1; (Yang, 1994). The Γ model for rate variation assigns significant probability near zero when the α < 1 (accommodating invariant sites). The Γ(2,1) hyperprior on α assigns 95% probability to the interval (0.36, 4.74), allowing for small and large values. Four independent runs for each gene set produced concordant topologies. We diagnosed MCMC performance using Tracer 1.7 (Rambaut *et al*., 2014).

### *Wolbachia* and mtDNA haplotyping of black and lined color morphs

To confirm that the same *Wolbachia* strain infects different *P. ignipectus* populations and color morphs, we amplified and Sanger sequenced five protein-coding *Wolbachia* genes *(coxA, hcpA, fbpA, ftsZ,* and *wsp*) in both directions (Eurofins Genomics LLC, Louisville, Kentucky)(see below, Supplemental Table 2). We also amplified and Sanger sequenced *gatb*, but sequence quality was consistently too low to include in our analyses. Samples included one infected female of each color form (black or lined), from each of the four populations (Carthage, New Portland, New Vineyard, and Strong) sampled in both years (Supplemental Table 1).

To specifically assess if *Wolbachia* might contribute to the morphological contact zone between New Vineyard (monomorphic black) and New Portland (monomorphic lined) *P. ignipectus*, we also we amplified and Sanger sequenced the *cytochrome C oxidase I* (*CoI*) mitochondrial locus from one male and one female from these populations, with the exception of one (New Vineyard black male) that did not produce usable sequence. We also produced *CoI* sequences for one black and one lined female from the polymorphic Strong population.

We visually inspected each sequence for quality and ambiguities, and consensus sequences were used as queries for a BLASTn search and the NCBI “nr” database to confirm that orthologous genes were amplified (Altschul *et al.*, 1990). We then used the “multiple locus query” function of the multi locus sequence typing (MLST) database to type *Wolbachia* (Baldo *et al*., 2006). Together these data enable us to test for differentiation in *Wolbachia* and mtDNA between populations and color forms, including between populations monomorphic for different color forms separated by only 10 km in Maine.

### Analysis of CI loci

Recent work has identified CI-causing factors *(cifs)* associated with WO prophage in *Wolbachia* genomes (Beckmann, Ronau and Hochstrasser, 2017; LePage *et al.*, 2017; Shropshire *et al.*, 2018; Shropshire and Bordenstein, 2019; Shropshire, Leigh and Bordenstein, 2020). Two genes (*cifA/B*) transgenically expressed in male *D. melanogaster* induce CI, while one gene *(cifA)* expressed in females rescues it. To identify *cif* loci, we used BLASTn to search for *cif* homologs in our whole genome raw reads, querying the Type 1 *cif* pair in *w*Mel, the Type 2 pair in *w*Ri, the Type 3 pair in *w*No, the Type 4 pair in *w*Pip, and the Type 5 pair in *w*Stri (Lindsey *et al*., 2018; Bing *et al*., 2020; Martinez *et al*., 2020). We later broadened our search for Type 1 pairs by querying *w*Pip and *w*NPa pairs (Klasson *et al*., 2008; Gerth and Bleidorn, 2017). For each Type, we extracted raw reads that covered at least 40% of the genes. We then corrected and assembled the reads with canu 2.1.1 (Koren *et al*., 2017, 2018; Nurk *et al*., 2020), producing sequences with about a 1% error rate. We limit our analyses to the discovery of *cif* types, since we did not generate additional sequence data to further correct the long reads. The assembled genes were compared to those in Martinez et al. (2020).

### Analysis of *Wolbachia* frequency variation

To test for *Wolbachia* frequency variation, we extracted DNA from many individuals from each collection using a standard squish buffer protocol and identified *Wolbachia* infections using polymerase chain reaction (PCR) (Simpliamp ThermoCycler; Applied Biosystems, Singapore) (Meany *et al*., 2019). We amplified the *Wolbachia* surface protein (*wsp*) (Braig *et al*., 1998) and arthropod-specific 28S rDNA, which served as a positive control (Baldo *et al*., 2006) (Supplemental Table 2). PCR products were visualized using 1% agarose gels. Assuming a binomial distribution, we estimated exact 95% confidence intervals for *Wolbachia* frequencies for each collection. We used Fisher’s exact test (FET) to determine differences in frequencies among sites, between years, between sexes, and between color forms.

## Results

### *P. ignipectus* likely acquired its group-B *Wolbachia* following initial divergence from *P. bicincta*

Across all samples, *Wolbachia* infection frequency (*p*) in *P. ignipectus* is high (*p* = 0.93 [0.90, 0.95]; *N* = 486). Based on five Sanger sequenced loci, the multiple sequence query of the MLST database supports that a group-B strain, most closely related to *Wolbachia* in Chloropidae (Diptera) (ID 93, ST 104), infects our *P. ignipectus* samples—we call this strain *w*Pig. Preliminary phylogenetic analyses using only our five Sanger sequenced genes also placed *w*Pig in group B. Our draft *w*Pig assembly size (1.32Mb, N50 = 91,011) falls in the range of complete *Wolbachia* genomes (e.g., *w*Mel at 1.26Mb and *w*Ri at 1.44Mb), despite its fragmentation (50 contigs). In total, we extracted 65 single-copy homologs of equal length (43,473 total bp) for our phylogenetic analysis, which also places *w*Pig in group B (Figure 2). When excluding the *w*Pig genome, we were able to extract an additional 135 homologs (16,7241 bp) from *w*Mel, *w*Ri, *w*Pip, and *w*Mau. This indicates that significant residual error in the *w*Pig assembly reduces the number of homologs meeting our equal length criteria for inclusion. Finer placement of *w*Pig among group-B strains will require the generation of short-read data to further correct our draft *w*Pig assembly. Thus, we do not attempt to place *w*Pig precisely among group-B strains.

**Figure 2.**
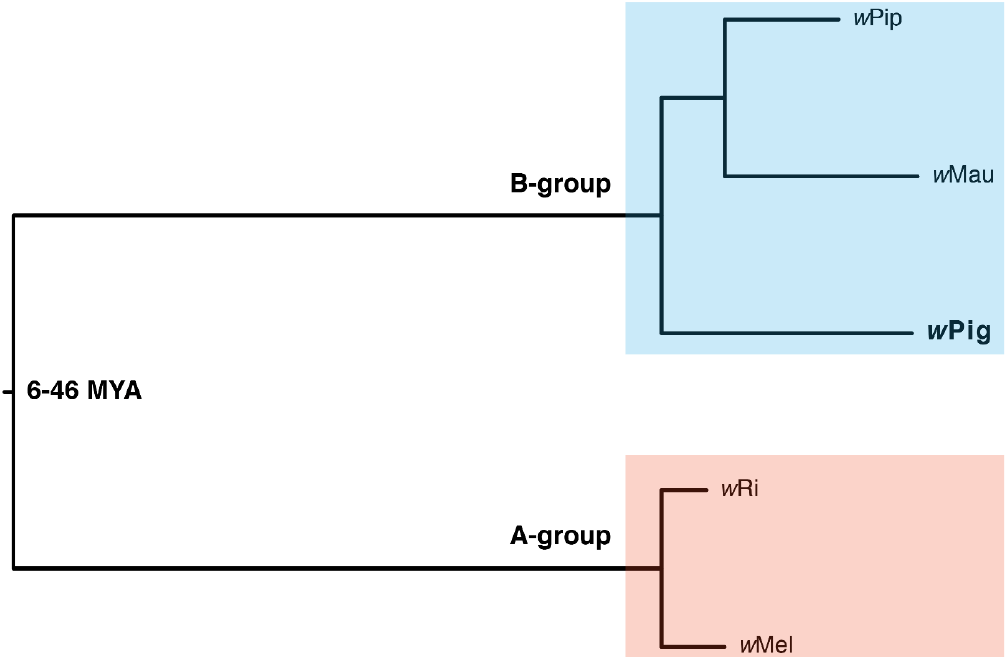
An estimated phylogram for model group-A (wRi, Klasson *et al*., 2009); and (*w*Mel, Wu *et al*., 2004) and group-B (*w*Pip_Pel, Klasson *et al*., 2008); and (*w*Mau, Meany *et al*., 2019) *Wolbachia,* plus *w*Pig. All nodes have Bayesian posterior probabilities of 1. The divergence time of groups A and B is superimposed from (Meany *et al*., 2019). The phylogram shows significant variation in the substitution rates across branches, with long branches separating groups A and B.

None of the *P. bicincta* samples from Hawaii and Florida were *Wolbachia* infected. Even if some *P. bicincta* are *Wolbachia* infected, as previously reported for one individual used as a PCR control in another study (Anderson, Rustin and Eremeeva, 2019), *Wolbachia* infection frequency (*p*) must be very low across the *P. bicincta* range, given our species estimate and credible interval *(p* = 0.0 [0.0, 0.04]; *N* = 100), keeping in mind the possibility that the Hawaiian population may have experienced a recent bottleneck during introduction and may not be representative of the species in the native range. Very low frequency *Wolbachia* infections in global *P. bicincta* populations, in combination with generally high *w*Pig frequencies in *P. ignipectus*, indicates that *P. ignipectus* likely acquired *w*Pig after its initial divergence from *P. bicincta*. Because testing predictions about modes of *Wolbachia* acquisition requires formal analysis of *Wolbachia*, host nuclear, and host mtDNA phylograms and chronograms, we are unable to distinguish between introgressive and horizontal *w*Pig transfer (Raychoudhury *et al.*, 2009; Conner *et al*., 2017; Gerth and Bleidorn, 2017; Turelli *et al*., 2018; Cooper *et al*., 2019). We discuss this further below.

Of the additional species we netted from Little bluestem, all *L. quadrangularis* were uninfected *(p* = 0.0 [0.0, 0.14]; *N* = 25), all *P. spumarius* were infected (*p* = 1.0 [0.48, 1.0]; *N* = 5), and only one *P. killa* individual was infected *(p* = 0.04 [0.001,0.21]; *N* = 24). Because *Wolbachia* that infect *P. spumarius* and *w*Pig in *P. ignipectus* are both at high frequency, we also typed the *Wolbachia* infecting *P. spumarius* to determine if a *w*Pig-like variant also infects this host species. The multiple sequence query in the MLST database supports that a different group-B strain, most closely related to the thrip species *Aptinothrips rufus* (ID 1945, ST 509) infects *P. spumarius*. Generating more sequence data will be required to resolve the phylogenetically relationships of these and other group-B strains, including *Wolbachia* in *P. spumarius* (Lis, Maryańska-Nadachowska and Kajtoch, 2015).

### No apparent effect of *w*Pig on the maintenance of the morphological *P. ignipectus* contact zone

The Strong, Carthage, and Dixfield *P. ignipectus* populations (Figure 3) were polymorphic for the black and lined forms (Figure 1, Supplemental Table 1), like three populations close to Rumford, Maine sampled in earlier work (Thompson and Carvalho, 2016). This set of mixed color-form populations runs roughly from Rumford northeast to Strong, but not to the sharp boundary dividing the monomorphic black New Vineyard population form the monomorphic lined New Portland population. It has the appearance of a hybrid zone, but one that does not reach the definitive boundary between the forms. The existence of distinct color forms both within and between the populations sampled facilitated investigation of the relationship, if any, between *Wolbachia* infection and patterns of color form occurrence.

**Figure 3.**
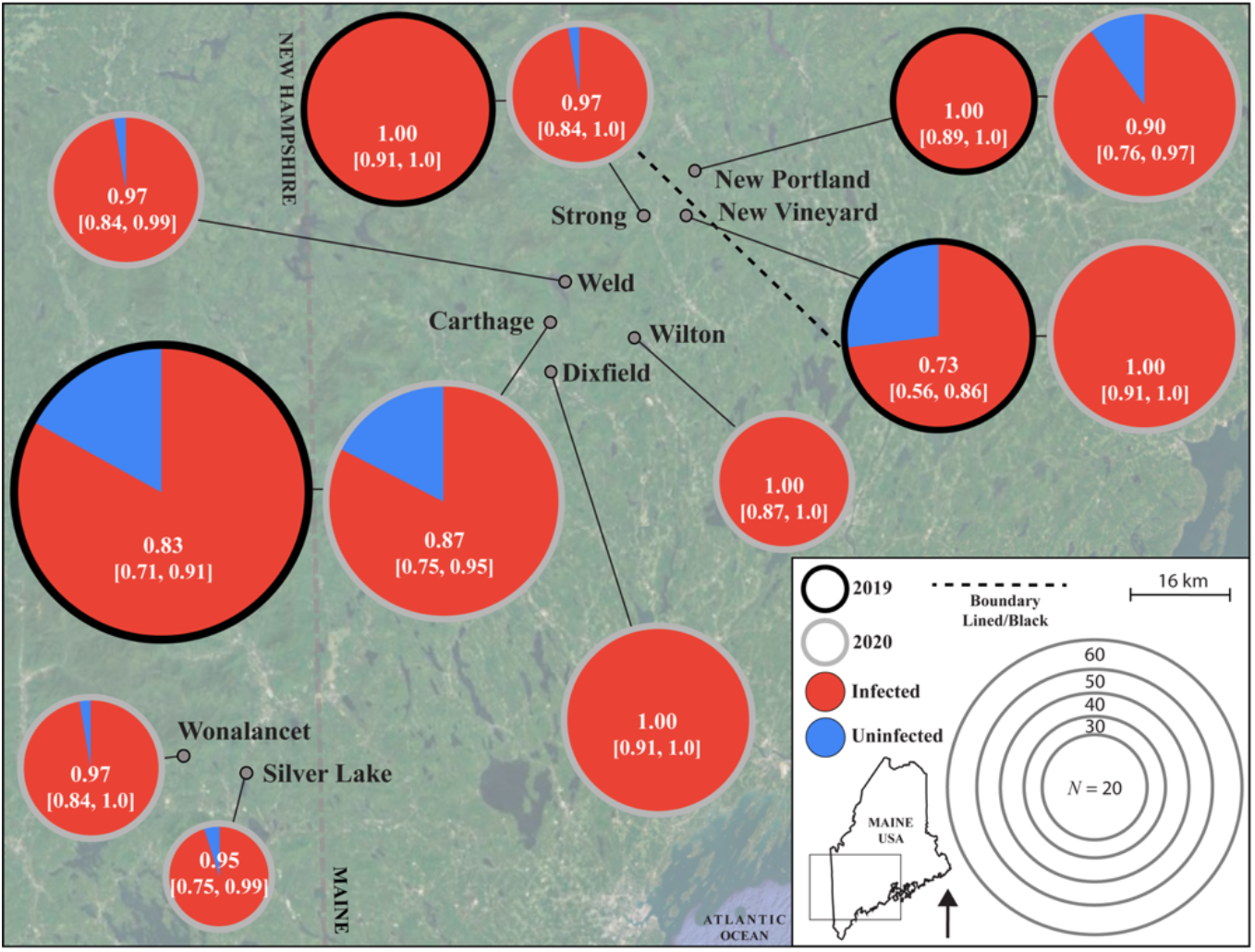
*w*Pig frequency varies through space and time. Circle size denotes sample size, with outline and fill color denoting sampling year and infection status, respectively. Sample means and 95% binomial confidence intervals are reported for each sample. The dashed back line denotes the geographical separation of monomorphic black and monomorphic lined *P. ignipectus* populations.

We found no evidence for *w*Pig genetic differentiation between *P. ignipectus* populations or color forms. Regions of the five *w*Pig genes we sequenced were identical, except for a single nucleotide position in *wsp*, where the Strong lined sample differed from all others. In addition to populations sharing *w*Pig type based on MLST loci, *w*Pig frequency did not vary between color forms (black: *p* = 0.93 [0.90, 0.95], *N* = 338; lined: *p* = 0.92 [0.86, 0.96], *N* = 123; FET, *P* = 0.69), among only males (black: *p* = 0.84 [0.75, 0.90], *N* = 98; lined: *p* = 0.90 [0.79, 0.97], *N* = 51; FET, *P* = 0.33), or among females (black: *p* = 0.97 [0.94, 0.99], *N* = 240; lined: *p* = 0.93 [0.85, 0.98], *N* = 72; FET, *P* = 0.19), across all samples. *w*Pig frequency also did not differ between New Vineyard (monomorphic black) and New Portland (monomorphic lined) populations (FET, *P* = 0.16).

We found no evidence for differentiation in *CoI* mtDNA haplotype between the New Vineyard and New Portland *P. ignipectus* populations, where all samples were identical across the 680 bp that we recovered. The black and lined females from the polymorphic Strong population also did not differ from each other, or from other populations, across this region. Thus, *w*Pig and mtDNA haplotypes were not differentiated between populations or color forms..

Our mtDNA haplotypes are also very similar to ten *P. ignipectus* samples included in the Barcode of Life Database (BOLD) (Foottit, Maw and Hebert, 2014). A single base-pair insertion present in all of our samples is absent from all ten BOLD samples. Four other sites in *CoI* that are polymorphic among the BOLD samples are fixed in our samples for one of the BOLD alleles. mtDNA haplotypes of *P. ignipectus* and *P. bicincta* also differ by less than 2 percent (Foottit, Maw and Hebert, 2014).

### The *w*Pig genome contains three divergent types of CI loci

We identified Type 1, 3, and 4 *cifs* in the *w*Pig genome (Martinez *et al*., 2020). This specific complement of *cifs* is not found in any other published *Wolbachia* genomes, but close relatives to each *w*Pig *cif* Type are. For instance, the *w*Pig Type 1 genes are 99% identical to those in the genome of the *Wolbachia* infecting the gall-inducing wasp *Diplolepis spinosa* (Cynipidae), but less than 90% similar to any others (Martinez *et al*., 2020). The Type 3 *w*Pig genes are 99% identical to those in the genome of the *Wolbachia* infecting *D. spinosa*, the Staphylinid beetle *Diploeciton nevermanni*, and the water strider *Gerris buenoi*. The *w*Pig Type 4 genes are 99% identical to those in *Wolbachia* infecting *Nomada* bees (*w*NLeu, *w*NFla, and *w*NPa), but less than 95% identical to other Type 4 *cifs*. The *Wolbachia* infecting *D. spinosa* does not have Type 4 *cifs*, distinguishing it from *w*Pig. None of the *w*Pig *cifs* are truncated relative to copies with 99% identity. Additional sequencing is required to make more detailed *cif* comparisons.

### Pervasive *w*Pig frequency variation

*w*Pig varied in frequency in several ways. First, frequency varied spatially among all samples (FET, *P* = 0.001)(Table 1), among sites in 2019 (FET, *P* < 0.0001), and 2020 (FET, *P* = 0.033). This variation occurred over a geographic radius of only 20 km in 2019 and 70 km in 2020 (Figure 3). Second, frequency varied across all samples between 2019 (*p* = 0.88 [0.82, 0.92]; *N* = 169) and 2020 (*p* = 0.95 [0.92, 0.97]; *N* = 317) (FET, *P* = 0.003). For the four sites we sampled in both years, frequencies were only significantly different between 2019 (*p* = 0.73 [0.56, 0.86]; *N* = 37) and 2020 (*p* = 1.0 [0.91, 1.0]; *N* = 40) in New Vineyard (FET, *P* < 0.001). Third, across all samples *w*Pig frequency was higher in females (*p* = 0.95 [0.93, 0.97]; *N* = 332) than males (*p* = 0.86 [0.80, 0.91]; *N* = 154) (FET, *P* = 0.001). However, this was driven mostly by a paucity of infected males in New Vineyard (males: *p* = 0.69 [0.50, 0.84], *N* = 32; females: *p* = 1.0 [0.92, 1.0], *N* = 45; FET, *P* < 0.0001), with no differences in *w*Pig frequency between males and females in other populations. *w*Pig frequency in males was relatively low in 2019 (*p* = 0.17 [0.02, 0.48]; *N* = 12), but fixed in 2020 (*p* = 1.0 [0.83, 1.0]; *N* = 20). We interpret these results as pervasive spatial, and rare temporal and sex-specific, variation in *w*Pig frequency.

**Table 1.** *w*Pig infection frequencies in *P. ignipectus* at each sampled site across both years.

| Site | GPS coordinates | N | Infected | p [Confidence Interval] |
| --- | --- | --- | --- | --- |
| Carthage | 44 36 44N, 70 28 10W | 116 | 98 | 0.84 [0.77, 0.91] |
| New Portland | 44 52 17N, 70 07 00W | 72 | 68 | 0.94 [0.86, 0.98] |
| New Vineyard | 44 45 14N, 70 08 01W | 77 | 67 | 0.87 [0.77, 0.94] |
| Strong | 44 47 08N, 70 13 42W | 69 | 68 | 0.99 [0.92, 1.0] |
| Silver Lake | 43 53 01N, 71 10 41W | 20 | 19 | 0.95 [0.75, 1.0] |
| Dixfield | 44 34 10N, 70 27 21W | 41 | 41 | 1.0 [0.91, 1.0] |
| Weld | 44 41 27N, 70 25 30W | 33 | 32 | 0.97 [0.84, 1.0] |
| Wilton | 44 37 58N, 70 18 10W | 26 | 26 | 1.0 [0.87, 1.0] |
| Wonalancet | 43 54 38N, 71 21 29W | 32 | 31 | 0.97 [0.84, 1.0] |

Sample sizes (*N*), infection frequencies (*p*), and exact 95% binomial confidence intervals for each site.

## Discussion

Our results suggest that *w*Pig is a group-B *Wolbachia* acquired after the initial divergence of *P. ignipectus* from *P. bicincta*. Analysis of *Wolbachia* and mtDNA haplotypes indicates that *w*Pig has no apparent effect on the *P. ignipectus* morphological contact zone in Maine. Across all samples, *w*Pig occurs at very high frequencies, consistent with our discovery of three divergent sets of CI loci in the *w*Pig genome. Finally, we document pervasive spatial, and rare temporal, *w*Pig frequency variation. We discuss this in more detail below.

### *Wolbachia* acquisition in spittlebugs

In contrast to very high *w*Pig frequencies in *P. ignipectus*, we found no evidence of *Wolbachia* in our sample of 100 *P. bicincta*. A prior report of one infected *P. bicincta* sample indicates that *Wolbachia* could infect this species (Anderson, Rustin and Eremeeva, 2019). If so, it must be at very low frequencies, given our credible interval here (*p* = 0.0 [0.0, 0.04]; *N* = 100). Mathematical models predict that intense CI drives *Wolbachia* to high frequencies, balanced by imperfect maternal transmission (Hoffmann, Turelli and Harshman, 1990; Turelli and Hoffmann, 1995); conversely, *Wolbachia* that do not cause strong CI tend to occur at much lower frequencies (Hamm *et al*., 2014; Kriesner *et al*., 2016; Cooper *et al*., 2017; Hague *et al*., 2020). While crossing to test for CI in the laboratory is not currently feasible in this system, the presence of three sets of CI loci in the *w*Pig genome, combined with its very high frequencies, suggests that *w*Pig causes intense CI.

How did *P. ignipectus* acquire *w*Pig? There are three possibilities: cladogenic transmission from its most recent common ancestor with its sister species, presumably *P. bicincta* or a close relative; by introgression from *P. bicincta* or another close relative; or by horizontal transmission (O’Neill *et al*., 1992). Given that we find no evidence for a high frequency *Wolbachia* in *P. bicincta*, cladogenic acquisition seems implausible. Without more extensive analysis of close relatives, we cannot rule out introgression. However, opportunities for introgression with species other than *P. bicincta* have likely been limited. Other species of the genus *Prosapia* or family Cercopidae occur no further north than the USA-Mexico border region, about 1,400 km from the nearest *P. ignipectus* populations and 3,000 km from the populations studied here.

Overall, the limited data are consistent with relatively recent non-cladogenic transmission, a process that seems to be common among *Drosophila* species (Turelli *et al.*, 2018). It may also be common among spittlebugs. This would be in stark contrast to obligate transovarial endosymbionts associated with amino acid nutrition in spittlebugs and other hemipterans (Koga *et al*., 2013). In addition to the thrip-related *Wolbachia* found in *P. spumarius* in this study, Nakabachi *et al.* (2020) report that two spittlebug species, *Aphrophora quadrinotata* Say and *Philaenus maghresignus* Drosopoulos & Remane (both *Aphrophoridae’),* harbor *Wolbachia* with 16S rRNA sequence that is identical to *Wolbachia* in two psyllid species, two whiteflies, an aphid, a planthopper, two leafhoppers, two grasshoppers, a mosquito and a weevil. Likewise, Lis *et al.* (2015) report that *Wolbachia* they studied in *P. spumarius* is closely related to strains in vespids, drosophilids, whiteflies, chrysomelid beetles and weevils based on five MLST loci. Kapantaidaki *et al.* (2021) also report *Wolbachia* infections at low levels in *P. spumarius,* as well as higher frequencies in *Neophilaenus campestris* (Fallén) (Aphrophoridae). Based on five MLST loci, their *N. campestris* strain is closely related to *Wolbachia* found in a leafhopper (Hemiptera) and cluster with *Wolbachia* from a planthopper, a scale insect and a psyllid (all Hemiptera), as well as two chrysomelid beetles, two butterflies, a parasitic wasp and a mosquito. Koga *et al.* (2013, table S3) report the presence of *Wolbachia* in the spittlebug *Cosmoscarta heros* (F.) (Cercopidae), in addition to *A. quadrinotata* and *P. maghresignus*.

In contrast, five specimens of *Poophilus costalis* (Walker) (Aphrophoridae) (Wiwatanaratanabutr, 2015), six specimens of *Philaenus tesselatus* Melichar (Lis, Maryańska- Nadachowska and Kajtoch, 2015), 37 specimens of *Philaenus signatus* Melichar (Lis, Maryańska-Nadachowska and Kajtoch, 2015; Kapantaidaki *et al*., 2021), and single specimens of *Philaenus arslani* Abdul-Nour & Lahoud, *Philaenus loukasi* Drosopoulos & Asche, and *Philaenus tarifa* Remane & Drosopoulos (Lis, Maryańska-Nadachowska and Kajtoch, 2015) were not infected. Based on limited sequence data, the emerging pattern suggests that *Wolbachia* infection is widespread, but far from ubiquitous among spittlebugs, and that when it does occur, it often involves *Wolbachia* strains similar to those infecting distantly related insects. Whole *Wolbachia* and host genomic data is sorely needed to test our hypothesis that horizontal *Wolbachia* acquisition might be common in spittlebugs.

### Little contribution of *w*Pig to the *P. ignipectus* morphological contact zone

We find no evidence for differentiation in *w*Pig or mtDNA haplotypes among *P. ignipectus* color forms. This includes the monomorphic black (New Vineyard) and lined (New Portland) populations that are separated by only 10 km in Maine, with no obvious barriers to dispersal or reproduction (Thompson and Carvalho, 2016). We also found no variation in *w*Pig or mtDNA haplotypes between black and lined individuals in the polymorphic Strong population. *w*Pig frequency also did not vary between color forms. These data indicate that *w*Pig is unlikely to significantly contribute to the maintenance of the *P. ignipectus* morphological contact zone.

How common are *Wolbachia* effects on host RI? Obligate *Wolbachia* infections in cooccurring *D. paulistorum* semi-species contribute to assortative mating and generate hybrid inviability and male sterility (Miller, Ehrman and Schneider, 2010). *Wolbachia* also contribute to reinforcement between *Wolbachia*-infected *D. recens* and uninfected *D. subquinaria* (Shoemaker, Katju and Jaenike, 1999; Jaenike *et al*., 2006). In contrast, *Wolbachia* do not contribute to premating, gametic, or postzygotic RI among the three *D. yakuba*-clade host species (Cooper *et al*., 2017). While the crossing schemes used in these *Drosophila* studies to dissect *Wolbachia* contributions to RI are not feasible in *P. ignipectus* and many other systems, our genetic data here lend support to our prior conjecture that *Wolbachia* contributions to RI observed in some *Drosophila* may be the exception rather than the rule (Turelli, Lipkowitz and Brandvain, 2014; Cooper *et al*., 2017).

### Pervasive *w*Pig frequency variation

Mathematical models indicate that imperfect maternal transmission, *Wolbachia* fitness effects, and the severity of CI govern *Wolbachia* frequencies in host populations. *Wolbachia* that cause intense CI tend to occur at high and stable frequencies, balanced by imperfect maternal transmission (Barton and Turelli, 2011; Turelli and Hoffmann, 1995; Hoffmann, Turelli and Harshman, 1990; Turelli and Hoffmann, 1991; Carrington *et al*., 2011; Kriesner *et al*., 2013); while *Wolbachia* that cause weak or no CI tend to persist at intermediate, often variable frequencies (Hamm *et al*., 2014; Kriesner *et al*., 2016; Cooper *et al*., 2017; Hague *et al*., 2020). Accumulating evidence for variable infection frequencies (Hamm *et al*., 2014; Kriesner *et al*., 2016; Schuler *et al*., 2016; Hughes *et al*., 2011; Lis, Maryańska-Nadachowska and Kajtoch, 2015; Cooper *et al.*, 2017), including our discovery here, highlights that infection frequencies are not static, even for high frequency variants.

With the exception of model systems like *w*Ri in *D. simulans*, few estimates of the key parameters required to approximate population frequency dynamics and equilibria of *Wolbachia* exist (Turelli and Hoffmann, 1995; Carrington *et al*., 2011). *w*Mel-like *Wolbachia* frequencies in the *D. yakuba* clade vary through space and time in west Africa (Cooper *et al*., 2017), due in part to effects of cold temperatures on *w*Yak titer (Hague *et al.*, 2020). CI strength also varies in the *D. yakuba* clade, which may influence infection frequencies (Cooper *et al.*, 2017; Hague, Caldwell and Cooper, 2020). *w*Mel frequencies vary with latitude in *D. melanogaster* populations, potentially due to *w*Mel fitness costs in the cold (Kriesner *et al.*, 2016). Interestingly, hot temperatures reduce *w*Mel CI strength and transmission in transinfected *Aedes aegypti* used for biocontrol of human disease (Ross *et al.*, 2017, 2020), suggesting that temperature may generally influence key parameters underlying *Wolbachia* infection frequencies.

What underlies variable *w*Pig frequencies in nature? High *w*Pig frequencies and the presence of three divergent sets of *cifs* suggest, but do not confirm, that *w*Pig causes strong CI. It seems plausible that some or all of these loci were horizontally acquired (Cooper *et al.*, 2019), but additional sequence data are required to test this. We hypothesize that variable *w*Pig transmission rates contribute to the frequency variation we observe, potentially due to environmental effects on titer, as observed for *w*Yak (Hague *et al*., 2020). Temporal variation in transmission was also observed for *w*Ri between two samples of *D. simulans* collected from Ivanhoe, California in April and November of 1993 (Turelli and Hoffmann, 1995; Carrington *et al.*, 2011), although the relative stability of *w*Ri frequencies in global *D. simulans* populations suggests that its transmission persists across a range of environmental conditions. Additional analyses of *Wolbachia* titer and transmission in the field, and across environmental contexts, are needed to better understand the causes of *Wolbachia* frequency variation. Comparing the titer and transmission of *Wolbachia* that occur at different frequencies in nature—for example, those that do and do not cause intense CI — would be particularly useful.

## Data Accessibility Statement

All data will be uploaded to DRYAD or GenBank upon acceptance.

## Author Contributions Section

**Timothy B. Wheeler:** Data curation, Investigation, Validation, Visualization, Writing - original draft, Writing - review & editing. **Vinton Thompson:** Conceptualization, Data curation, Formal analysis, Investigation, Methodology, Project administration, Resources, Visualization, Writing - original draft, Writing - review & editing. **William R. Conner:** Data curation, Formal analysis, Investigation, Writing - original draft, Writing - review & editing. **Brandon S. Cooper:** Conceptualization, Data curation, Formal analysis, Funding acquisition, Investigation, Methodology, Project administration, Resources, Supervision, Validation, Visualization, Writing - original draft, Writing - review & editing.

## Acknowledgments

We thank M. Thorne and A.G. Dale for *P. bicincta* collections and D. McVicar and F. Selchin for access to the Wonaloncet site. Michael Turelli provided comments that greatly improved an earlier draft. We also thank M. Hague, D. Shropshire, and K. Van Vaerenberghe for very helpful comments. Research reported in this publication was supported by the National Institute of General Medical Sciences of the National Institutes of Health (NIH) under award number R35GM124701 to B.S.C., and by the University of Montana Genomics Core.

---

**Supplemental Table 1.**
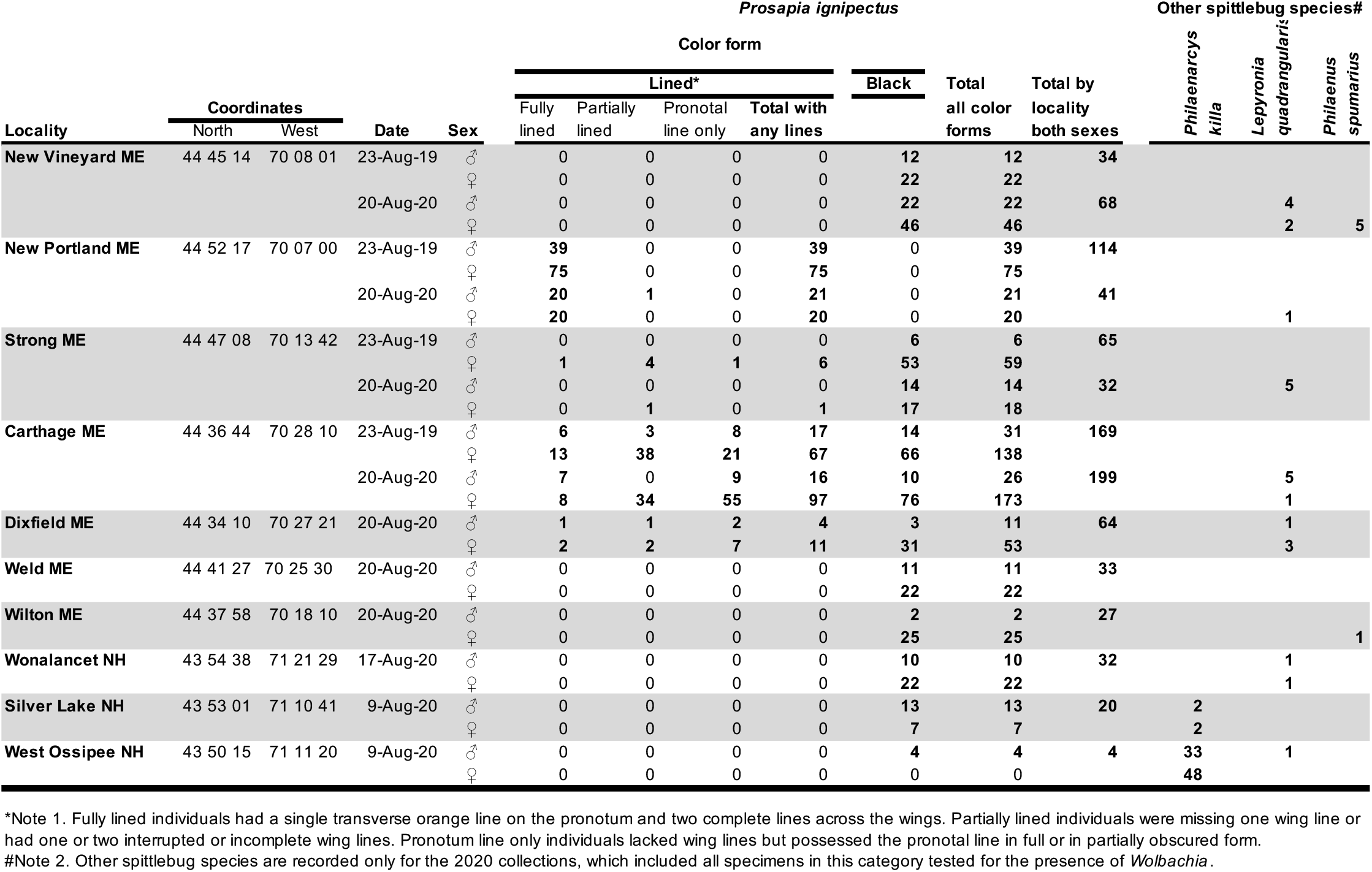
Spittlebugs from *Schizachyrium scoparium,* Maine and New Hampshire, August 2019 and 2020

**Supplemental Table 2.**
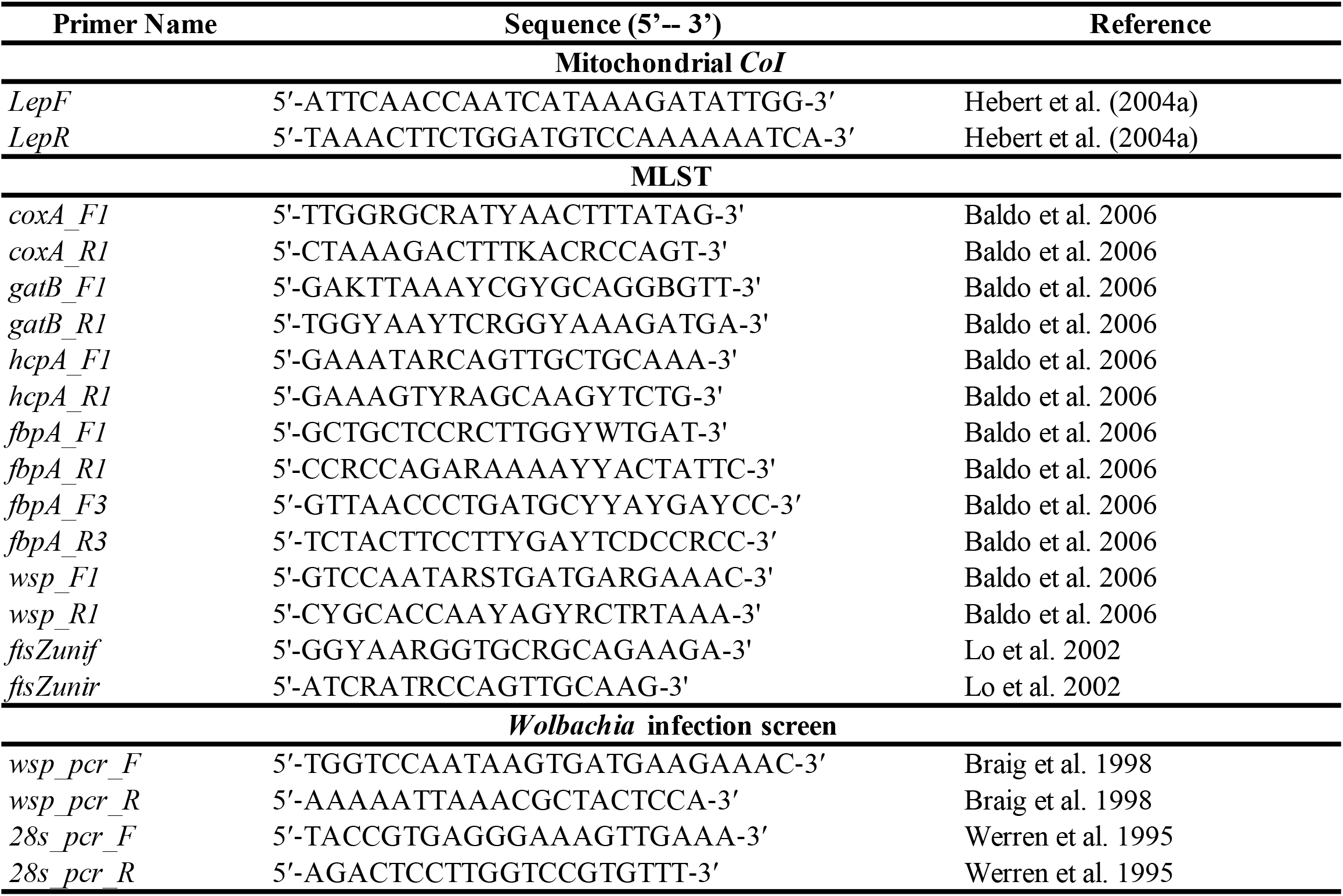
PCR primers used in this study

